# Diversity of the woodland strawberry inflorescences results from heterochrony antagonistically regulated by *FvTFL1* and *FvFT1*

**DOI:** 10.1101/2022.11.09.515873

**Authors:** Sergei Lembinen, Mikolaj Cieslak, Teng Zhang, Kathryn Mackenzie, Paula Elomaa, Przemyslaw Prusinkiewicz, Timo Hytönen

## Abstract

A vast variety of inflorescence architectures have formed during angiosperm evolution. Here we analyze the diversity and development of the woodland strawberry inflorescence. We show that it is a thyrse: a compound inflorescence in which the primary monopodial axis supports lateral sympodial branches, thus combining features of racemes and cymes. We demonstrate that this architecture is related to differences in the size and shape of the primary and lateral inflorescence meristems. We further show that woodland strawberry homologs of TERMINAL FLOWER 1 (TFL1) and FLOWERING LOCUS T (FT) antagonistically regulate the development of both the racemose and cymose components of the strawberry thyrse: the loss of functional *FvTFL1* and overexpression of *FvFT1* reduce the number and complexity of the cymose components, whereas silencing of *FvFT1* has the opposite effect and can partially rescue the *fvtfl1* mutation. We complement our experimental findings with a computational model, which captures the development of the woodland strawberry inflorescence using a small set of rules, and shows that its phenotypic diversity can be explained in terms of heterochrony resulting from the opposite action of FvTFL1 and FvFT1 on the progression from the branching to flowering state.

## INTRODUCTION

The arrangement of individual flowers in time and space is critical for plants’ reproductive success (Harder et al., 2004; Harder & Prusinkiwicz, 2013), and ultimately for agricultural yield and fruit uniformity (Eshed & Lippman, 2019). Many angiosperms organize their flowers into clusters called inflorescences. The structural arrangement of inflorescences has long attracted the attention of plant scientists, horticulturalists, and agronomists, who described, explained, and enhanced through breeding the diverse inflorescence architectures.

Based on their branching patterns, a vast variety of inflorescence architectures can be classified as monopodial or sympodial (Weberling, 1989; Prusinkiewicz & Lindenmayer 1990). In monopodial inflorescences, or racemes (Figure 1A), new flowers form at the lateral positions on the indeterminate or determinate primary axis. In contrast, in cymose or sympodial inflorescences (Figure 1B), new flowers form at the terminal positions, and the new growth axes are established laterally. The classification of inflorescence architectures is complicated by the occurrence of compound inflorescences. For instance, panicles, or polypodial inflorescences, consist of repetitively branching determinate or indeterminate monopodial axes (Figure 1C). In contrast, in thyrses, the primary indeterminate or determinate (Prenner et al., 2009) monopodial axis bears sympodial branches (Figure 1C).

**Figure 1.**
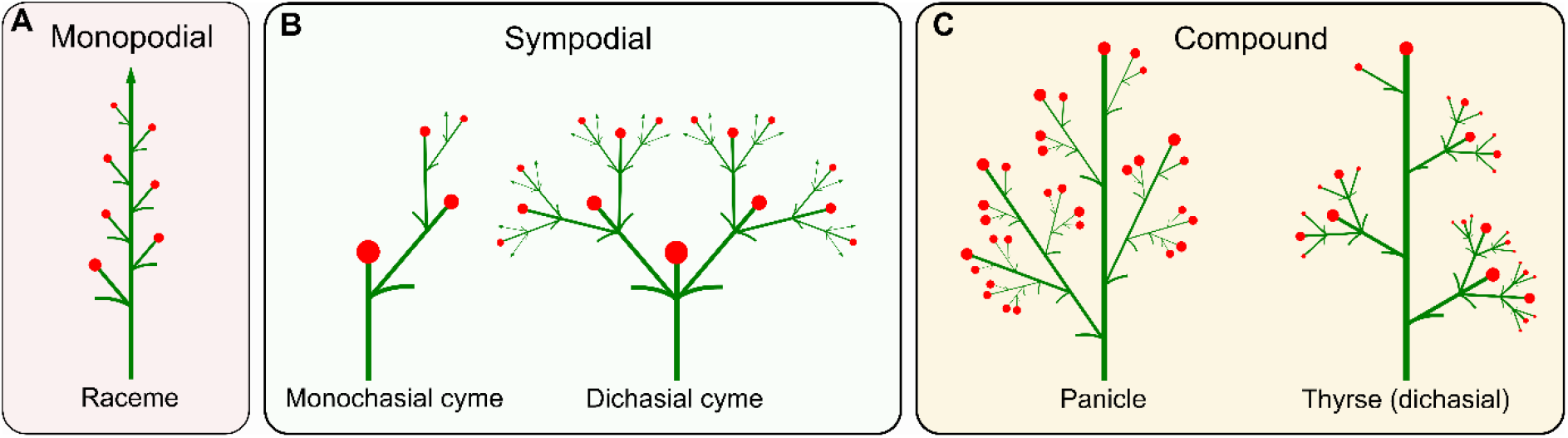
Inflorescence branching types. **A**. Monopodial branching of a simple raceme. **B**. Sympodial branching of monochasial and dichasial cymes. **C**. Compound branching of a panicle and a thyrse (dichasial)

The development of inflorescence branching patterns is regulated by the activity of meristems, the small groups of multipotent cells located at the branch tips. Inflorescence meristems (IMs) are capable of initiating new IMs (i.e., branching axes) and eventually transition to flower meristems (FMs), which produce flower organs. Differences in the timing of these transitions – a form of heterochrony – control the diversity of inflorescence architectures (Grimes, 1993; Lemmon et al., 2016; Prusinkiewicz et al., 2007).

The molecular processes that regulate developmental decisions in meristems have been unraveled in model plants with monopodial as well as sympodial inflorescences. In simple racemes, such as in *Arabidopsis thaliana*, the IM state is maintained by TERMINAL FLOWER 1 (TFL1). Plants lacking a functional TFL1, which is normally expressed in the center of the developing IM, produce determinate inflorescences with a few flowers (Alvarez et al., 1992; Schultz & Haughn, 1993). On the IM flanks, the expression of *LEAFY* (*LFY*) and *APETALA1* (*AP1*) activates the floral organ identity genes, promoting FM identity. Inactivation of *LFY* prevents specification of FM identity and transforms flowers into shoots (Weigel et al., 1992; Weigel & Meyerowitz, 1993).

Subsequent studies in Arabidopsis revealed that a *TFL1* homolog *FLOWERING LOCUS T* (*FT*) is also involved in regulating inflorescence architecture (Lee et al., 2019). FT and TFL1 belong to a family of phosphatidylethanolamine-binding proteins (PEBPs) and integrate environmental signals such as photoperiod and temperature to control transition to reproductive growth (reviewed in McGarry & Ayre 2012; Rantanen et al., 2015). FT is a mobile protein which is produced in leaves and moves into the shoot apical meristem (SAM) to promote flowering (Corbesier et al., 2007). In racemose Arabidopsis, the loss of functional FT promotes meristem indeterminacy and inflorescence branching (Lee et al., 2019). Further experiments revealed that FT and TFL1 proteins compete for the same binding partner, a bZIP transcription factor FD, and form complexes via 14-3-3 proteins (Zhu et al., 2021). When formed, FT-FD and TFL1-FD complexes act as transcriptional activators and repressors, respectively, affecting multiple target genes including the meristem identity regulators *LFY* and *AP1* (Zhu et al., 2020).

Studies in plants with cymose inflorescences have revealed both similarities and differences in the genetic control of inflorescence development, compared to racemes. For example, in petunia (*Petunia hybrida*) and tomato (*Solanum lycopersicum*), both members of the *Solanaceae* family, the orthologs of *LFY* — *FALSIFLORA* (*FA*) and *ABERRANT LEAVES AND FLOWERS* (*ALF*) — are expressed in the terminal meristems and promote the transition to FM state (Molinero-Rosales et al., 1999; Souer et al., 2008). However, to fully establish FM identity, FA and ALF require a co-factor. In petunia, DOUBLE TOP (DOT), an ortholog of UNUSUAL FLOWER ORGANS (UFO), physically interacts with ALF to establish the FM identity (Souer et al., 2008). Interestingly, inactivation of *SELF-PRUNING* (*SP*), an ortholog of *TFL1* in tomato, does not directly affect inflorescence architecture (Pnueli et al., 1998). In contrast, it was shown that *FT* homolog *SINGLE FLOWER TRUSS* (*SFT*; Molinero-Roslaes et al., 2004) regulates inflorescence branching in tomato (Park et al., 2012; Quinet et al., 2006).

Woodland strawberry (*Fragaria vesca* L.) is a perennial rosette plant and a model for the octoploid cultivated strawberry (*Fragaria* × *ananassa*) and the Rosaceae family in general (Edger et al. 2018). Historically, inflorescence architecture of *Fragaria* has been classified as pleiochasial cyme (Valleau, 1918), dichasial cyme (Anderson & Guttridge 1982; Guttridge, 1985), cyme (Menzel, 2019) and corymb (Gleason & Cronquist, 1991). This diversity of terms reflects the extensive morphological variability of strawberry inflorescences, which was highlighted almost a century ago (Darrow, 1929). The variability was attributed to the environment, genetic differences between specific cultivars (Darrow, 1929; Foster & Janick 1969; Anderson & Guttridge 1982), or to the sex of the flowers (Ashman & Hitchens, 2000). However, the developmental and genetic basis of inflorescence architecture in strawberry has never been elucidated.

Here we analyze inflorescence architecture and development in woodland strawberry and begin to dissect the underlying molecular mechanism regulating its development. We show that the inflorescences of woodland strawberry have compound architecture, combining a monopodial primary axis with sympodial lateral branches. We summarize these findings in a computational model of thyrse development, which supports the diversity of strawberry inflorescence architectures due to heterochrony controlled by strawberry homologs of *FT* and *TFL1*.

## RESULTS

### Inflorescence of woodland strawberry is a determinate thyrse

To understand the variability of strawberry inflorescences, we analyzed inflorescence development in diploid woodland strawberry. A woodland strawberry inflorescence has a monopodial primary axis terminated by a flower. This axis typically supports two (Figure 2A) and occasionally three (Figure 2B) secondary branches separated by long internodes. In many monopodial structures, the main axis is relatively straight and can be recognized easily. In woodland strawberry, however, secondary branches often assume a dominant position, continuing approximately in the direction of their supporting internodes, whereas the primary axis changes direction at each branching point. To identify the course of the primary axis we thus relied on the positions of bracts (b; Figure 2) associated with the branches they subtend.

**Figure 2.**
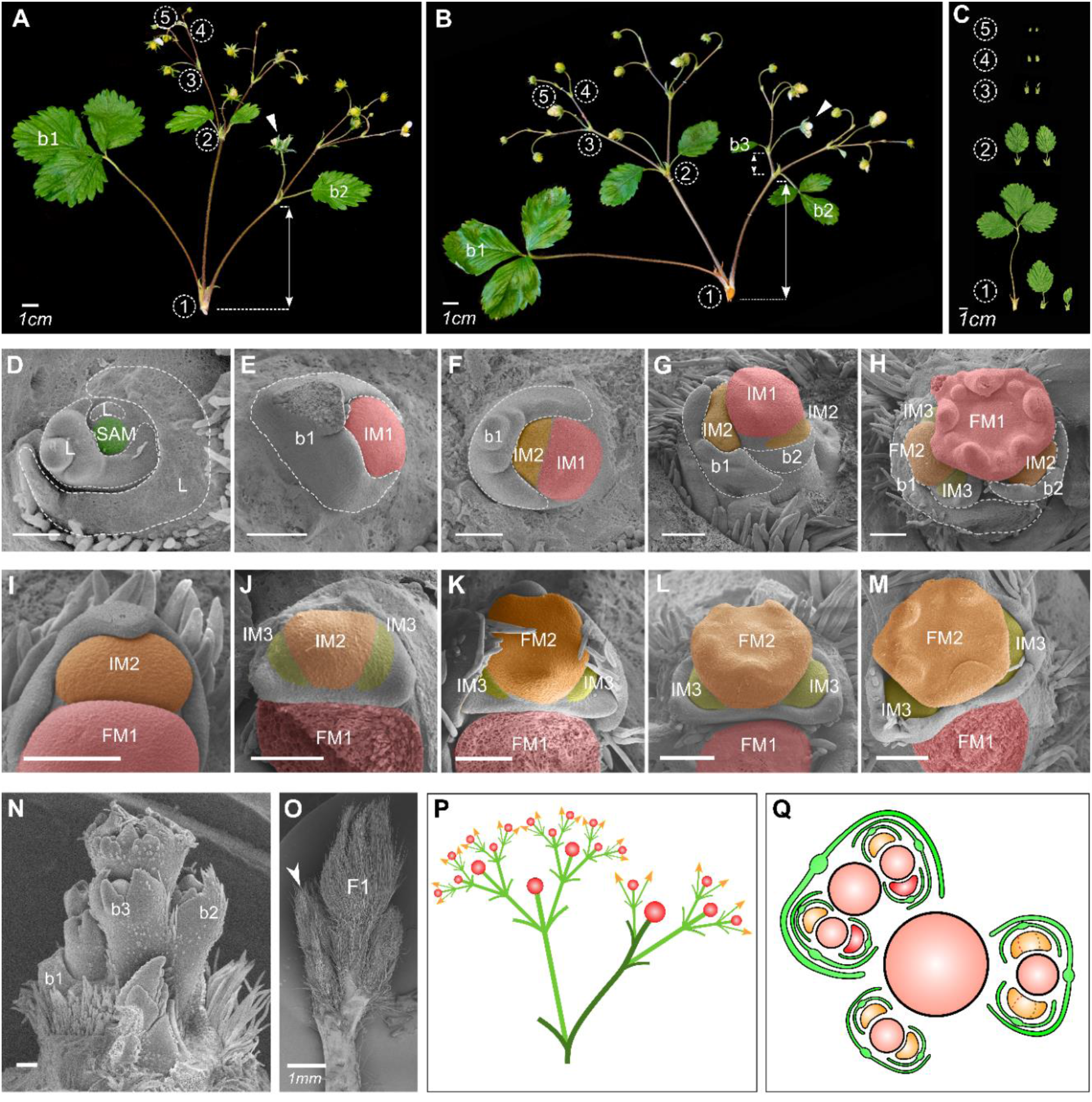
Architecture and development of thyrse inflorescence in woodland strawberry. **A – B**. Inflorescence architectures of wild-type woodland strawberry plants with two **(A)** and three **(B)** lateral branches on the primary axis. White arrowheads indicate the primary flowers; white lines with arrows indicate the distances between bracts on the primary branching axes; numbers in circles indicate branching iterations **C**. Typical bract phenotypes at branching points of different orders. **D–H**. Development of the primary inflorescence meristem (IM). **D**. Round shoot apical meristem (SAM) with monopodially produced leaves. **E**. Early IM1 of increased size, compared to the SAM. **F**. Initiation of the first lateral IM2. **G**. Initiation of the second lateral IM2. **H**. IM1 transitioned into flower meristem (FM1). **I–M**. Development of the lateral meristems. **I**. Elongated, crescent-like shape of IM2. **J**. An almost simultaneous initiation of the lateral IM3s at the ends of IM2. **K–M**. Transition of IM2 into FM2 and development of IM3s. **N**. Primary axis with three bracts, each subtending a lateral meristem. **O**. The first lateral branch (white arrow) displaces the primary flower (F1) and assumes a more dominant position. **P**. Schematic representation of thyrse inflorescence with leading lateral branch. The primary axis is highlighted in darker green. **Q**. Schematic representation of relative meristem positions during the development of woodland strawberry thyrse. Relatively more confined IMs are highlighted in red. Scale bars in **D–N** equal 100 µm. b – bract.

The architecture of lateral branches is different from that of the main axis. Each secondary axis supports a pair of third-order branches subtended by bracts, then terminates with a secondary flower. The third-order branches are approximately opposite each other: the internode between them is practically absent. This branching pattern repeats with each third-order branch producing a pair of fourth-order branches and a terminal flower, and typically continues up to fifth- or sixth-order branches. At high branching orders, only one lateral branch may emerge, although both bracts are present (Jahn & Dana 1970). In spite of this departure from symmetry, the structures supported by the primary axis are best characterized as dichasial cymes (Figure 1B; Weberling, 1989; Prenner et al., 2009). With the determinate primary monopodial axis supporting sympodial branches, inflorescence of woodland strawberry qualifies as a determinate thyrse (Figure 1C).

Characterizing these inflorescences further, we observed that they are basitonic, i.e., the branches originating near the inflorescence base are more elaborate than those near the top (Lück et al., 1990). The first lateral branch growing from the axil of the leaf-like bract b1 is larger and bears more flowers than the branch that originates from the axil of b2 (Figure 2, A and B). The third lateral branch, if present, is even smaller and produces fewer flowers than the second or the first lateral branch. The internode between b2 and b3 is also shorter than between b1 and b2 (Figure 2, A and B).

Moreover, the size of successive bracts is dramatically reduced, and the shape is simplified from a three-lobed structure resembling a leaf to a bracteole-like structure with a single lobe (Figure 2C). Within branches, the bracts are more similar in shape, but also decrease in size with each successive branching iteration.

### Two meristem types produce determinate thyrse of woodland strawberry

To investigate the ontogenesis of the strawberry inflorescence, we obtained a developmental sequence of inflorescence meristems using SEM imaging (Figure 2, D−M). The transition to flowering is associated with bulging and an almost two-fold increase in the size of the primary inflorescence meristem (i.e., the meristem that generates the primary inflorescence axis: IM1), compared to the vegetative shoot apical meristem (SAM) (Figure 2, D and E). IM1 has an approximately circular cross-section and produces bracts and lateral meristems (IM2) that are almost as large as IM1. The first bract (b1) is initiated during the SAM to IM1 transition (Figure 2E), with a lateral meristem (IM2) then forming in the bract axil (Figure 2F). As the primary meristem (IM1) continues to grow, an additional one or two bracts with associated lateral meristems (IM2) arise sequentially (Figure 2, G, H and N), leading to the monopodial architecture of the main axis. Due to their size, these bracts and meristems push the primary meristem (IM1) to the side (Figure 2O), which changes the course of the main axis at each branching point (Figure 2P) as evident in the mature inflorescences (e.g., Figure 2, A and B). The patterning of the main axis is terminated by the primary meristem acquiring flower identity and switching to the production of flower organs (FM1; Figure 2H).

The second and higher-order inflorescence meristems extend along the perimeter of their parent meristems and have an elongated, crescent-like shape (Figure 2, F and G). In consequence, the space needed to form the next-order meristems with associated bracts emerges concurrently near both ends of their parent meristem (Figure 2, I and J). These primordia become meristems that may produce the next iteration of the same pattern, while the parent meristem continues to grow without changing direction and develops into a flower (Figure 2, K–M). The compound dichasial structure of the woodland strawberry inflorescence thus results.

However, one end of each meristem of the third order (IM3) is inevitably closer to the inflorescence center than the other end, which leads to the uneven restriction of space available for the development of the fourth order primordia (IM4; Figure 2Q). A similar asymmetry occurs in higher order meristems, which may be the cause of, or contribute to, the commonly observed departure from symmetry at high branching orders, where only one lateral branch is present.

### FvTFL1 controls inflorescence architecture in woodland strawberry

In many plant species, inflorescence architecture and flowering time are tightly connected (Lee et al., 2019; Lifschitz et al., 2006; Liu et al., 2013). With this in mind, we looked for the genetic regulators that can affect architecture and development of the strawberry thyrse. In woodland strawberry, a mutation in *FvTFL1* accelerates flowering and reverses the photoperiodic requirement for flower induction from short-day to long-day (Koskela et al., 2012). To understand whether FvTFL1 also controls inflorescence development in woodland strawberry, we analyzed the inflorescence architecture and branching in seven *fvtfl1* mutants and nine wild type (WT) genotypes collected across Europe (Supplemental Figure S1, A–C). All *fvtfl1* cultivars used in this study were found to have a 2-bp deletion in the first exon of *FvTFL1* (Supplemental Figure S2) putatively leading to a nonfunctional FvTFL1 protein.

We found that the number of flowers per inflorescence was reduced more than two-fold in the *fvtfl1* mutant plants. On average, WT plants produced 14.2 ± 1.2 flowers per inflorescence, while *fvtfl1* mutants produced only 6.5 ± 1.4 flowers per inflorescence (p < 0.0001; Figure 3A). This reduction of flower numbers was associated with changes in the general architecture of inflorescences, particularly with dramatic changes in the number of lateral branches on the primary axis (Figure 3B−D). In plants with functional FvTFL1, we observed a high proportion of inflorescences with two (72%), and three (27%), lateral branches produced on the primary axis. Only 1% of the inflorescences had a single secondary branch. In contrast, the majority of inflorescences of *fvtfl1* mutant plants (78%) produced only one secondary branch (Figure 3C) and the remaining 22% had two secondary branches. Such a dramatic change in the proportion of lateral branches supported by the primary axis suggests that in *fvtfl1* mutants IM1 transitions to FM1 before it is able to produce the second lateral branch on the primary axis.

**Figure 3.**
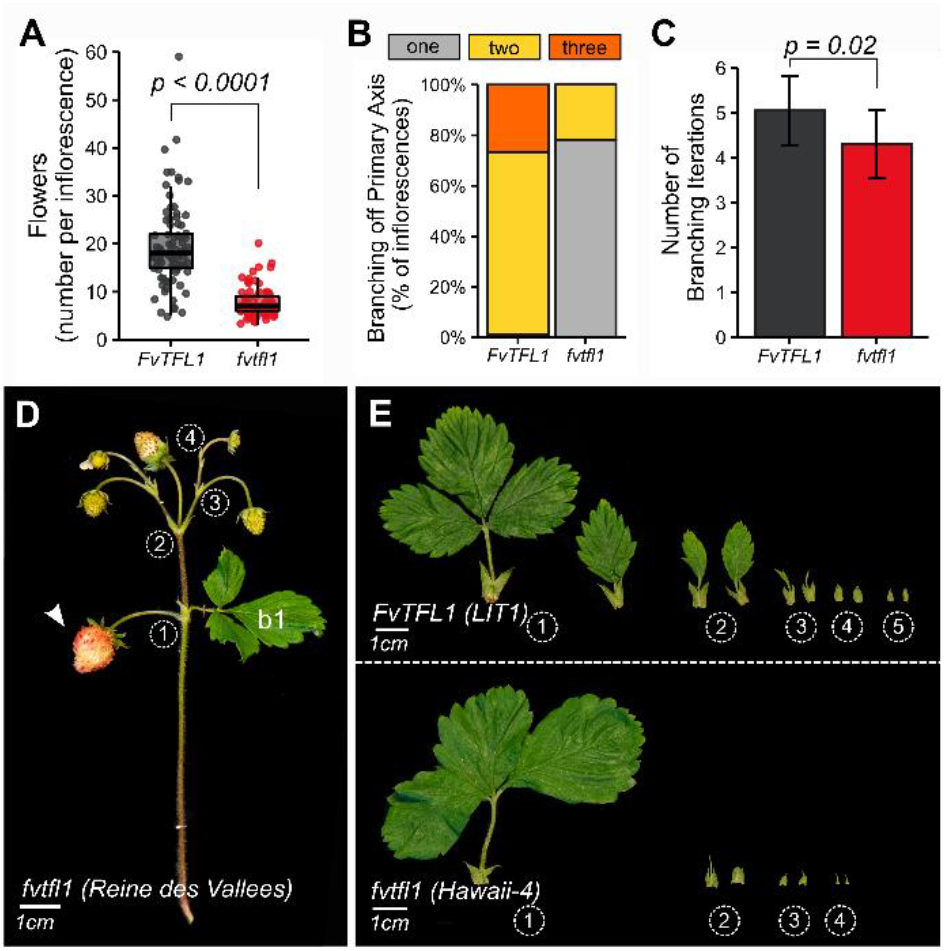
FvTFL1 controls inflorescence architecture in woodland strawberry. **A**. Number of flowers on the first inflorescence of seven *fvtfl1* (red) and nine *FvTFL1* (grey) genotypes. Boxplots and points show the distribution of raw data. Each point represents an individual inflorescence. **B**. Percentage of observed inflorescences with one, two, or three branches on the primary branching axis. **C**. Average number of branching iterations along the longest branching path. Bars and whiskers represent the mean ± standard deviation. Data in **A** and **C** were analyzed using a generalized linear mixed model (GLMM), with accessions and cultivars nested within *fvtfl1* and *FvTFL1* groups. P values for significant differences between *fvtfl1* and *FvTFL1* groups of plants are shown (*t-*test). **D**. Inflorescence phenotype of *fvtfl1* cultivar Reine des Vallées; white arrow indicates the position of the primary flower. b – bract. **E**. Typical bract phenotypes along the longest branching path of *FvTFL1* and *fvtfl1* genotypes. Numbers in circles indicate branching iterations.

In the lateral cymose branches, the loss of functional FvTFL1 did not affect the dichasial branching pattern. However, the total number of branching iterations in the *fvtfl1* mutants (4.3 ± 1.1) was reduced compared to the WT plants (5.1 ± 1.0; p = 0.02; Figure 3D). To understand the reasons behind this variation we compared the bracts of *fvtfl1* mutants and WT plants (Figure 3E). The first bract (b1) on the primary axis was the largest and typically had three lobes in plants with functional and non-functional FvTFL1. Furthermore, in both groups of plants, each successive lateral IM produced smaller bracts; however, at each corresponding branching point, WT plants had larger bracts than *fvtfl1* mutants.

### *FvFT1* regulates inflorescence architecture in *fvtfl1* background

Next, we decided to investigate the role of *FvFT1* in the regulation of inflorescence architecture. Previously, FvFT1 was found to promote flowering under long-day conditions in the *fvtfl1* mutant background (Rantanen et al., 2014; Kurokura et al., 2017). We analyzed the inflorescence architecture and branching in *fvtfl1* plants (*Hawaii-4*) with silenced (RNAi) and overexpressed (35S) *FvFT1* (Supplemental Figure S3). Overexpression (OX) of *FvFT1* caused a pleiotropic effect by dramatically reducing the overall plant and leaf size and quickly transforming all shoots into inflorescences (Supplemental Figure S4). *Hawaii-4* plants typically produced inflorescences with 5.7 ± 1.9 flowers, whereas transgenic plants with constitutive *FvFT1* expression produced inflorescences with only 1 – 3 flowers (p < 0.0001; Figure 4, A, D and E). In contrast, silencing of *FvFT1* caused an almost two-fold increase in the number of flowers per inflorescence (12 ± 0.24; p = 0.006; Figure 4A). The main reason for the higher number of flowers per inflorescence was the formation of the second lateral branch along the primary axis, as in the genotypes with functional FvTFL1. We observed that about 80% of the inflorescences in *FvFT1-RNAi* plants initiated two lateral branches on the primary axis, compared to only 20% in *Hawaii-4* plants (Figure 4B). Furthermore, *FvFT1-RNAi* plants typically produced more branching iterations than *Hawaii-4* (p = 0.0001; Figure 4, C, D and F). The branching of the primary axis in *fvtfl1* mutant plants (*Hawaii-4*) with silenced *FvFT1* expression was similar to WT plants with functional FvTFL1, except that we did not observe inflorescences with three lateral branches on the primary axis in *FvFT1-RNAi* lines (Figure 4F). Moreover, we found that *FvFT1-RNAi* plants occasionally produced an intermediate phenotype, where the second bract (b2) on the primary axis was associated with the sepals of the primary flower (Figure 4G). A similar phenotype was observed in the *FvFT1-OX* lines, however in these lines the first bract (b1) became associated with the sepals of the primary flower (Figure 4H).

**Figure 4.**
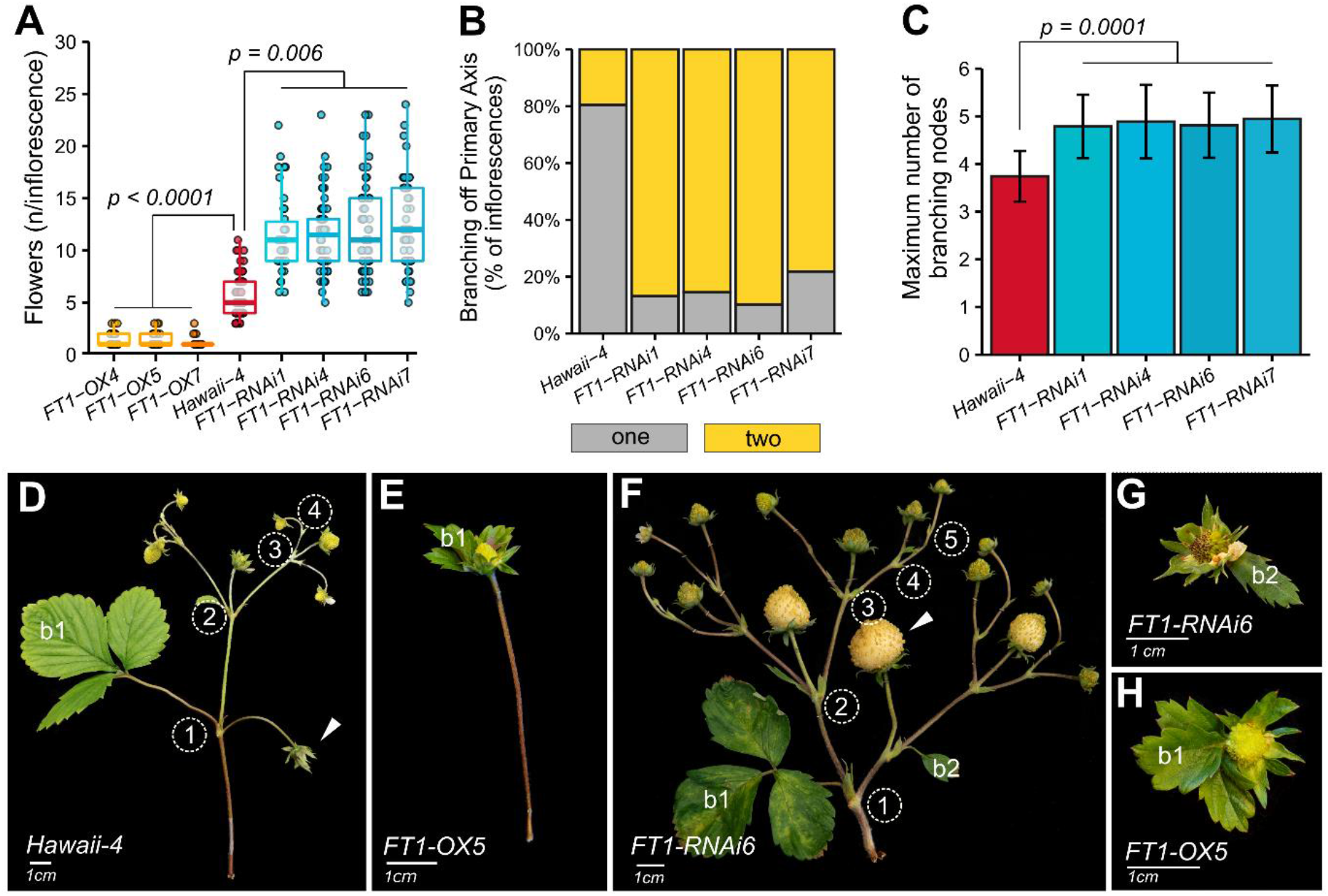
FvFT1 regulates inflorescence architecture in woodland strawberry. **A**. Number of flowers per inflorescence in *FvFT1* overexpression *(FT1-OX), FvFT1* silenced (*FT1-RNAi*), and control *(Hawaii-4)* plants. Boxplots and points show the distribution of raw data. Each point represents an individual inflorescence. Up to five inflorescences per plant were examined. **B**. Maximum number of branching iterations in *FvFT1-RNAi* lines and control *(Hawaii-4)* plants. Bars and whiskers show the mean ± standard deviation of the raw data. Data in **A** and **B** were analyzed by fitting a generalized linear mixed model (GLMM); plants were grouped within construct type (OX, RNAi, or control). P values for significant differences between OX/RNAi and control are shown (*t*-test). **C**. Percentage of inflorescences with one and two branches along the primary axis. **D–F**. Inflorescence phenotypes of control *(Hawaii-4), FT1-OX5* and *FT1-RNAi6* plants. The numbers in circles indicate branching iterations. **G–H**. Intermediate phenotypes of *FvFT1-RNAi* and *FvFT1-OX* plants associating the second (b2) or first (b1) bract of the primary axis with the sepals of the primary flower. b – bract.

In the plants with the functional FvTFL1, the overexpression of *FvFT1* did not cause as severe a reduction in the overall plant size (Supplemental Figure S5A) as in *FvFT1-OX//fvtfl1* plants (Supplemental Figure S4). The number of flowers per inflorescence of *FvFT1-OX* plants with functional FvTFL1 background was reduced compared to WT and *fvtfl1* mutant plants (Supplemental Figure S3B). However, it was higher than in the *FvFT1-OX* plants with *fvtfl1* background. Altogether our data thus suggests the direct antagonistic functions of FvFT1 and FvTFL1 in inflorescence development of woodland strawberry.

### FvTFL1 and FvFT1 antagonistically regulate the inflorescence architecture in woodland strawberry

To understand the mechanism of the second lateral branch formation on the primary axis we analyzed WT (*FIN56*), *fvtfl1* (*Hawaii-4*) and *fvtfl1//FvFT1-RNAi* plants using SEM imaging (Figure 5, A–C). We compared the WT and *fvtfl1* plants at a stage when the primary FMs showed a similar developmental phase. The first lateral meristem (IM2) of *FIN56* with functional FvTFL1 was at a later developmental stage (Figure 5A) than the first (and single) lateral meristem (IM2) of *Hawaii-4* (Figure 5B). In *FIN56*, the lateral IM2 already formed a pair of axillary meristems (IM3s) and sepal primordia, while in *Hawaii-4*, only the bracts could be clearly distinguished on the single lateral IM2. A similar difference was also evident when *FvFT1* was silenced in *fvtfl1* background (Figure 5C). Overall, these findings suggest that the primary IM1 of the plants with functional FvTFL1 or silenced *FvFT1* develops into FM1 slower than in *fvtfl1* mutants, thus allowing formation of additional bracts and lateral IM2s.

**Figure 5.**
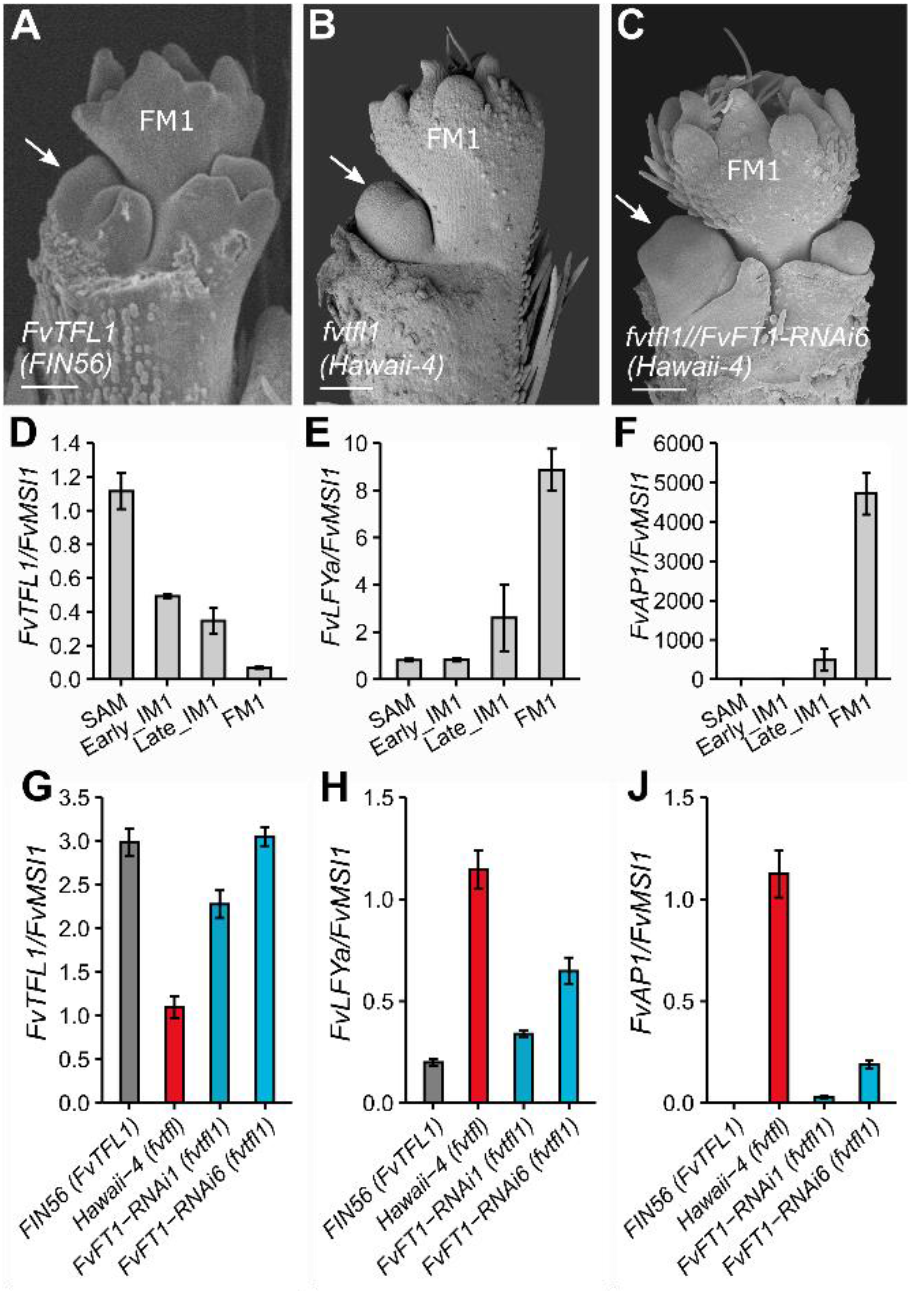
FvTFL1 and FvFT1 antagonistically shape the inflorescence of woodland strawberry. **A–C**. Primary axis development in *FvTFL1 WT* **(A)**, *fvtfl1* mutant **(B)**, and *FvFT1-RNAi* **(C)** plants. White arrows denote the first lateral IM2. Scale bars equal 100µm. **D–F**. The expression pattern of *FvTFL1* **(D)**, *FvLFYa* **(E)**, *and FvAP1* **(F)** in different meristems of *FvTFL1 WT* (*FIN56*) during transition to flowering. **G–J**. The expression pattern of *FvTFL1* **(G)**, *FvLFYa* **(H)**, and *FvAP1* **(J)** in the SAMs of FIN56 (WT), *Hawaii-4* (*fvtfl1*) and *FvFT1-RNAi* (*Hawaii-4*) plants. Bars and whiskers show the mean ± standard error (n = 3 – 5). Expression levels were normalized to SAM (D –F) or *Hawaii-4* (G–J). *FvMSI1* was used as a calibrator gene.

In woodland strawberry, *FvTFL1* is highly expressed in the vegetative shoot apex and gradually downregulated during flowering induction (Koskela et al. 2012). Molecular antagonism between *TFL1* and *LFY* has been described as the mechanism controlling inflorescence architecture in Arabidopsis (Schultz & Haughn, 1993). Subsequent studies in other species found that *AP1* may substitute *LFY* as *TFL1* antagonist in inflorescence development (Kobayashi et al., 2012; Ma et al., 2017). Here we observed that the decrease of *FvTFL1* expression is associated with the progression of the vegetative SAM towards FM identity (Figure 5D). The expression of the strawberry homologs of *LFY* and *AP1* showed the opposite pattern (Figure 5, E and F). The expression of both *FvLFYa* (*FvH4_5g09660*) and *FvAP1* (*FvH4_6g29600*) began increasing in late IM1s, where *FvTFL1* was still present. *FvTFL1* had the lowest expression in the FM tissues, concordant with the highest expression of *FvLFYa* and *FvAP1*. Altogether, our data suggests that the role of TFL1 in maintaining IM identity (Alvarez et al., 1992; Schultz & Haughn, 1993) is conserved in woodland strawberry.

To further elucidate the mechanism by which FvFT1 regulates inflorescence architecture in the absence of functional FvTFL1, we analyzed the expression levels of *FvTFL1, FvLFYa* and *FvAP1* in the SAMs of *FIN56* (*FvTFL1*), *Hawaii-4* (*fvtfl1*) and *FvFT1-RNAi* (*fvtfl1*) plants. We observed that the expression of *FvLFYa* and *FvAP1* was lower in the SAMs of *FIN56* and *FvFT1-RNAi* plants compared to *fvtfl1* (*Hawaii-4*) (Figure 5, H and J). In addition, the expression of *FvTFL1* was the lowest in the *fvtfl1* mutant (Figure 5G). Our results thus suggest that FvTFL1 and FvFT1 antagonistically control the transition to FM identity by regulating the expression of *FvLFYa* and *FvAP1*.

### A computational model supports heterochrony as the key determinant of the strawberry inflorescence diversity

We constructed a parametrized computational model of the woodland strawberry inflorescence to show that the observed diversity of the inflorescence architecture can be attributed to the regulation of the rate of development by *FvTFL1* and *FvFT1*. The model is expressed as an L-system, with separate rules capturing the monopodial development of the primary axis and the sympodial development of branches. Generic rules for both types of branching (c.f. Prusinkiewicz & Lindenmayer, 1990) are extended with a mechanism that controls transitions of individual meristems to flowers and terminates the formation of new meristems at the end of inflorescence development. These processes are controlled by a synthetic variable (integrating many influences) called *vegetativeness* (*veg*) (Prusinkiewicz et al., 2007), which is related to the concepts of a controller of phase switching (Schultz and Haughn, 1993) and meristem maturation (Park et al., 2012). In the strawberry model, *veg* decreases monotonically with time (Figure 6A). Its values are compared to two thresholds, *th*_*m*_ and *th*_*s*_, which are pertinent to the fate of monopodial and sympodial meristems, respectively. As long as *veg ≥ th*_*m*_, the primary meristem periodically produces lateral primordia subtended by bracts (Event 1 in Figure 6B^**\***^). This process terminates when *veg* drops below *th*_*m*_, at which point the meristem produces a terminal flower (Event 2). A lateral primordium develops in turn into a lateral meristem supported by a bract (Event 3), which subsequently produces two next-order primordia and a terminal flower (Event 4). This process periodically iterates as long as *veg* ≥ *th*_*s*_, giving rise to a compound dichasial cyme. Eventually *veg* drops below *th*_*s*_, which results in the production of bracts without associated meristems (Event 5) and arrests further branching. The distinct development of branches originating at the same node, in particular the case when only one lateral branch is present, are captured by introducing a small difference in the value of threshold *th*_*s*_ between sibling meristems. This difference may be related to the unequal space available for the development of primordia on opposite sides of an elongated meristem (Figure 2Q).

**Figure 6.**
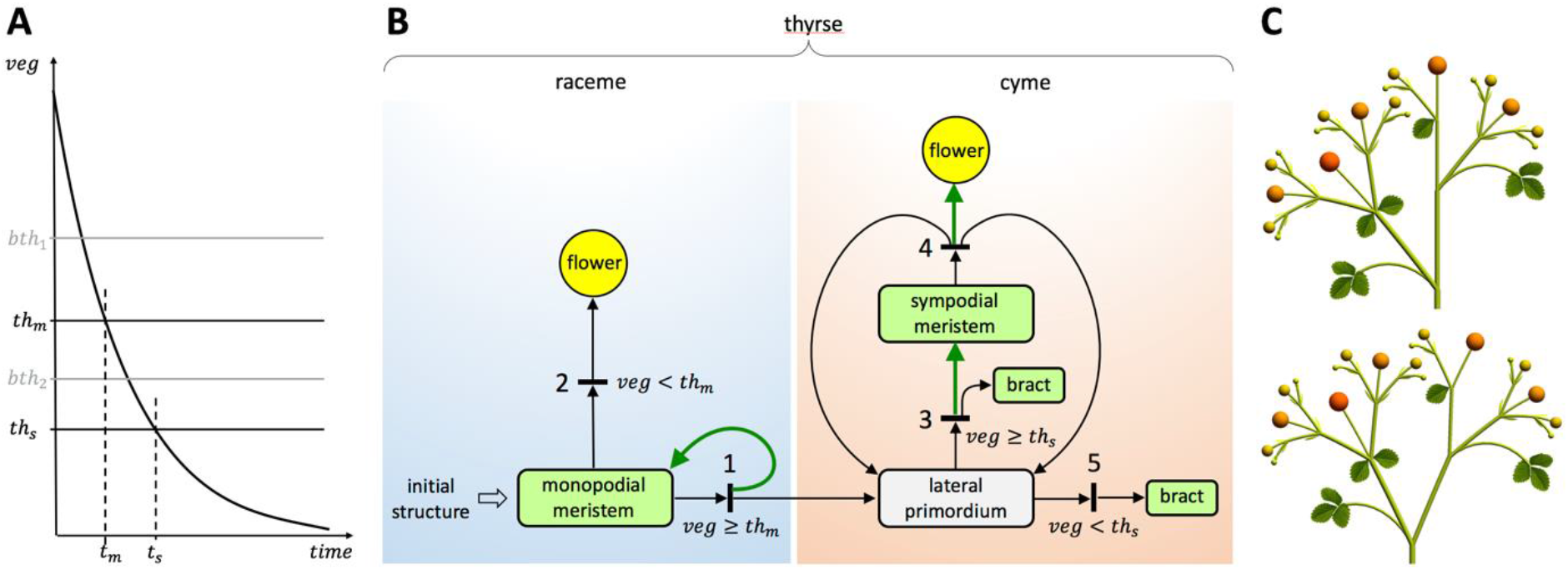
Developmental model of a woodland strawberry thyrse. **A**. Control of a thyrse development by *vegetativeness* (*veg*). Crossing threshold *th*_*m*_ terminates the production of primordia by the primary (monopodial) meristem; crossing threshold *th*_*s*_ terminates production of lateral primordia by the sympodial meristems. Times *th*_*m*_ and *th*_*s*_ at which these thresholds are crossed determine the inflorescence complexity (extent of branching). Additional thresholds, *bth*1 and *bth*_*2*_, control the transitions of bracts from 3-lobed to 1-lobed to narrow-scale forms. **B**. Petri net representation of the meristem production and fate. Circles and rectangles represent plant organs. The term ‘lateral primordium’ denotes an incipient structure yielding a lateral meristem supported by a bract, or a bract alone. Short black bars represent events taking place during development. These events are labelled by a number and are associated with conditions under which they may take place. Arcs with arrows represent relations between these events and the resulting structures. Two arrows originating in the same rectangle indicate alternative organ fates. Two or three arrows emanating from the same bar indicate production of multiple organs. Green-colored arrows indicate the introduction of an internode. **C**. The impact of the monopodial axis course on thyrse geometry. In the top model the monopodial axis is straight; in the bottom it deviates in the direction opposite to the lateral branch, as commonly observed in woodland strawberry (c.f. Figure 2 A, B, O and P).

In addition to the thresholds controlling branching architecture, the model includes thresholds *bth*_*1*_ and *bth*_*2*_ (Figure 6A), which control the transition of bracts from the 3-lobed form to 1-lobed form and to narrow scales, respectively. The model also includes a number of parameters and growth functions that control the geometry of the phenotypes of interest. Parameter values and functions can be chosen such that the simulated inflorescence development (Supplemental Movie 1) closely approximates observations (Figure 7). In particular, the deviation of the primary axis from a straight course substantially affects the overall appearance of the inflorescence, obscuring its thyrsoid architecture (Figure 6C). For details see the supplemental Model Code, Model Description, and the parameter values listed in Supplemental Table S2.

**Figure 7.**
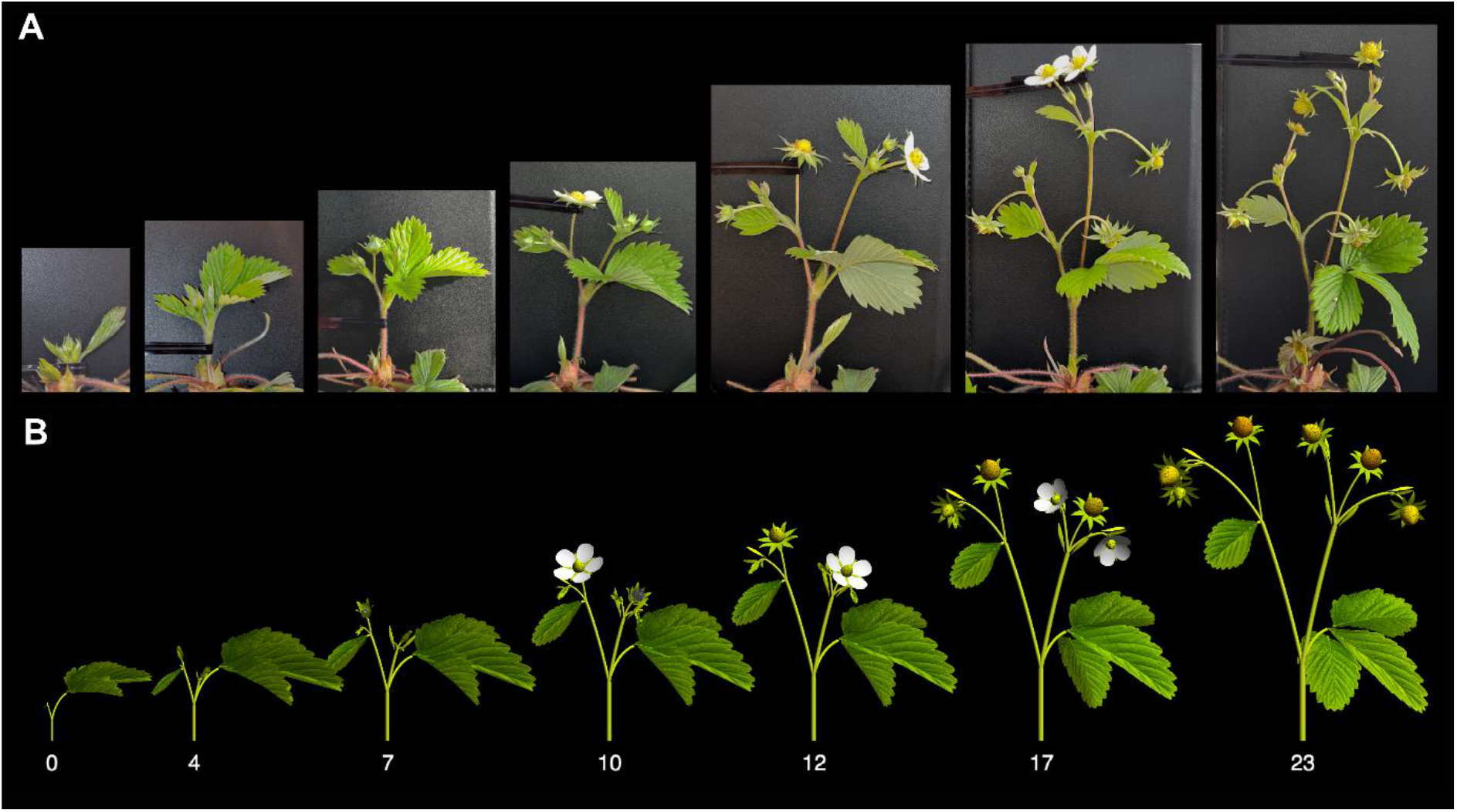
Development of a woodland strawberry inflorescence. **A**. Snapshots of inflorescence growth in a WT (GER12 x FIN2) woodland strawberry. Photographs were taken with the inflorescence supported in a vertical position to expose the branching pattern. Plants were induced to flower under 11°C and a 12h/12h (day/night) photoperiod for 6 weeks, then moved to 17°C and a 18h/6h photoperiod during inflorescence growth. Numbers indicate days of observation. **B**. Simulated development calibrated to sequence (A).

We experimented with the model to capture the key inflorescence features of the plants with different constitutions of *FvTFL1* and *FvFT1*. The most prominent trait that distinguishes the architectures collected in Figure 8A is the decreasing complexity (maximum order) of branching, caused by an accelerated cessation of branching. This decrease is accompanied by the accelerated progression of bracts from the three-lobed leaf-like form at the most basal position to the one-lobed and small-scale forms at more distal positions. Both the branching architecture and bract form are controlled in the model by the initial value and rate of decline of *veg*, and the thresholds *th*_*m*_, *th*_*s*_, *bth*_*1*_, and *bth*_*2*_ with respect to *veg* (Figure 6A). We observed that the sequence of inflorescence architectures (branching topologies) presented in Figure 8A can be reproduced simply by gradually increasing the rate of decline of *veg*, with minimal adjustments to other parameters except for the extreme phenotype of the *fvtfl1//FvFT1-OX* plants (Figure 8B; Supplemental Movies 2–8; Supplemental Table S1). A similar effect is caused by gradually reducing the initial value of *veg*. This result supports the conclusion that the diversity of strawberry inflorescences is a manifestation of heterochrony, in which *FvTFL1* decelerates, and *FvFT1* accelerates the rate of *veg* decline, respectively.

**Figure 8.**
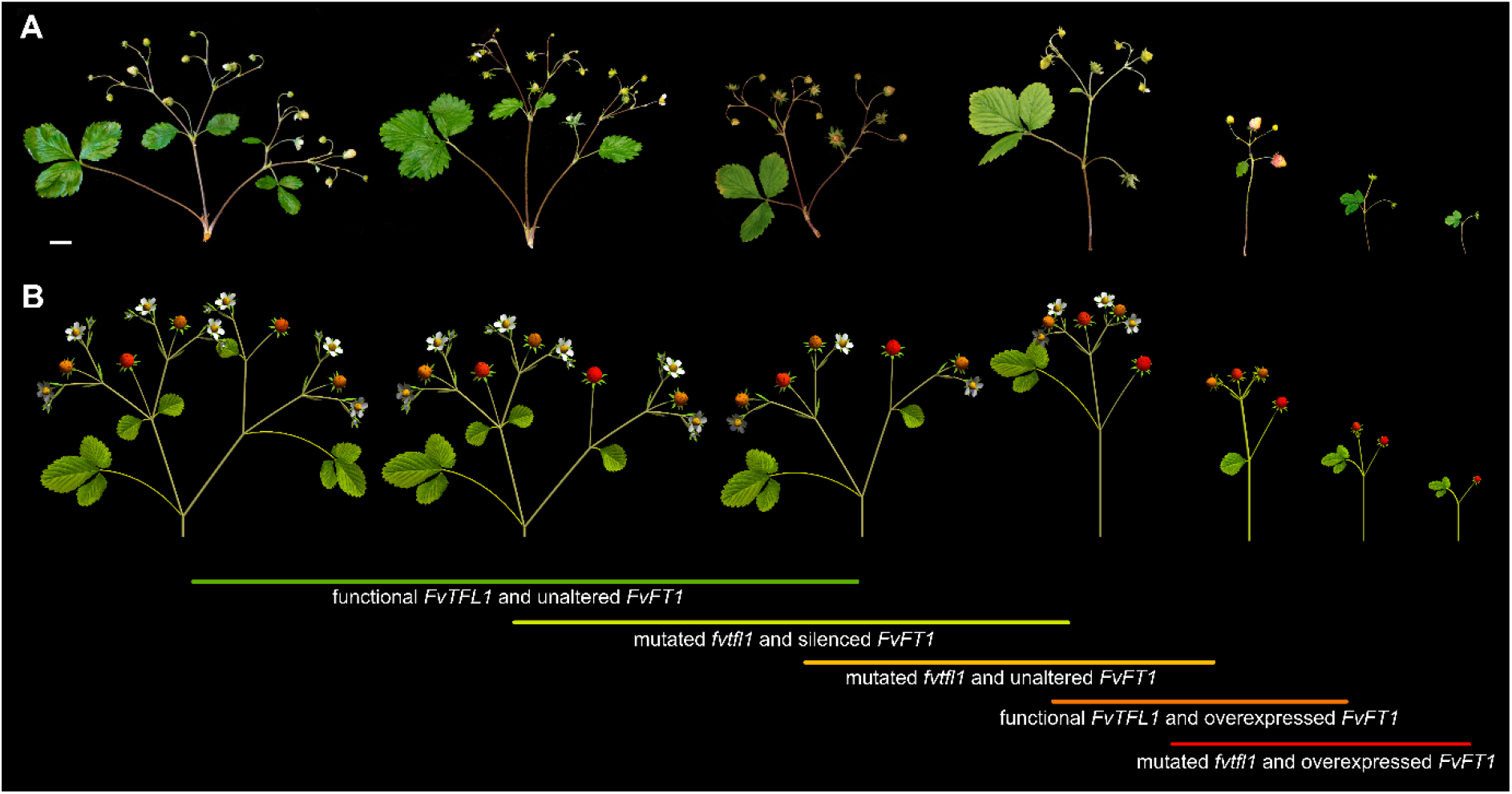
Diversity of woodland strawberry inflorescence phenotypes can be reproduced by altering model parameters. **A**. Observed diversity of inflorescence phenotypes. Scale bar equals 1 cm. **B**. Phenotypes reproduced by the model. Lines with text describe the genetic constitution of the plants. The color of the line indicates the rate of *veg* decline (green – slower, red – faster).

## DISCUSSION

Almost a century ago, Darrow (1929) pointed out the extensive variability of inflorescence architectures in cultivated strawberry *(F*. × *ananassa)* varieties, which he attributed to genetic differences between cultivars, environmental factors, and the vigor of individual plants. As outlined in Darrow’s inflorescence diagrams, the variation in inflorescence architecture of strawberries is mainly due to the variable number of branches on the primary inflorescence axis and branching iterations in the lateral branches. Here, using diploid woodland strawberry, we established the developmental and genetic basis of this variation. We have shown that the inflorescences of woodland strawberry consist of a monopodial determinate primary axis supporting sympodial lateral branches, an architecture characterized as a determinate thyrse (Prenner et al., 2009). This architecture arises from differences between the primary and lateral inflorescence meristems.

We have further demonstrated that the number of lateral branches on the primary monopodial axis and the number of iterations in the lateral sympodial branches are oppositely controlled by the two PEBP family proteins FvTFL1 and FvFT1. These proteins act by changing the timing of developmental decisions that transition inflorescence meristems to the flowering state, and eventually terminate the formation of further meristems. We supported this observation with a computational model which captures the diversity of studied phenotypes.

### Two types of IMs produce thyrse inflorescence in woodland strawberry

In the ontogenic view of inflorescence diversity, distinct inflorescence architectures are produced by different types of meristems (Claßen-Bockhoff & Bull-Hereñu, 2013). Our data shows that the determinate thyrse of woodland strawberry, which combines monopodial and sympodial branching patterns, is produced by two types of IMs. The monopodial primary axis is produced by a primary IM (IM1), which originates from the SAM that has expanded and begun producing leaf-like bracts upon entering the reproductive phase. Like the SAM, IM1 has a circular cross-section and produces bracts acropetally. After producing one to three bracts, the primary IM acquires FM identity, thus forming a closed monopodial axis. Each of the bracts on the monopodial axis subtends the newly formed and rapidly growing lateral IM, which, unlike the primary IM, has a crescent-shaped cross-section. According to the first available space theory (Hofmeister, 1868), new organ primordia are initiated in the largest available space between the previous ones. The initially elongated shape of the lateral IMs thus promotes a dichasial branching pattern, by allowing the pairwise initiation of next-order bracts and branches near opposite ends of the meristem.

We observed that, despite their morphological differences, both types of IMs followed a similar developmental progression towards FMs. Moreover, along the monopodial axis and with each successive branching iteration, the leaf-like bracts and the internodes were gradually reduced. By associating this reduction with the monotonic decrease of *veg*, properly choosing thresholds *th*_*m*_ and *th*_*s*_ at which the activity of each type of IM terminates, and setting additional parameters (e.g., branching angles) to agree with the observation of reference plants, we have been able to model the development of the woodland strawberry thyrse. The model confirms the usefulness of *veg* as a controller of development and sets the stage for understanding the diversity of strawberry inflorescences.

### *FvTFL1* and *FvFT1* antagonistically regulate the complexity of woodland strawberry inflorescences

Previous studies uncovered the role of *FvTFL1* in the regulation of flowering time in woodland strawberry (Koskela et al., 2012; Koskela et al., 2017). Flowering in WT strawberry requires downregulation of *FvTFL1*, which triggers the expression of the FM identity gene *FvAP1* (Koskela et al., 2012). Our results further show that *FvTFL1* is gradually downregulated as the SAM progresses towards the IM and then to the FM state. This expression pattern is different from that previously described in racemose and cymose plants. In monopodial Arabidopsis, *TFL1* is expressed in the SAM and IM and is specifically required to maintain IM identity (Alvarez et al., 1992; Schultz & Haughn, 1993). In sympodial tomato, *SP -* the closest homolog of *TFL1* - is expressed only in vegetative axillary meristems (Pnueli et al., 1998; Thouet et al., 2012). Concordantly, the *tfl1* mutation in Arabidopsis leads to the formation of a terminal flower on the otherwise indeterminate IM, whereas *sp* mutants in tomato do not show alterations of inflorescence architecture. The determinate primary IM of woodland strawberry is related to Arabidopsis in the sense that the loss of functional *FvTFL1* accelerates the formation of the terminal flower and cessation of the formation of lateral meristems. As a result, only one lateral branch is formed on the primary monopodial inflorescence axis of the *fvtfl1* mutant. Furthermore, in the lateral sympodial branches, the maximum number of branching iterations is reduced in the *fvtfl1* mutant plants, consistent with the accelerated termination of IMs.

Mutations in *FT* homologs were shown to affect the architecture in both monopodial and sympodial inflorescences. In Arabidopsis, FT, and its close homolog TSF, promote FM identity transition and act antagonistically to TFL1 (Lee et al., 2019). Furthermore, mutation in *SFT* was found to promote inflorescence branching in tomato (Park et al., 2012), suggesting a similar function. In woodland strawberry, FvFT1 was previously shown to promote flowering in *fvtfl1* mutant background (Koskela et al., 2012; Rantanen et al., 2014). Here we found that the role of *FvFT1* in inflorescence development of woodland strawberry is antagonistic to that of *FvTFL1*. Our data suggests that the faster progression towards FM in the *fvtfl1* mutant plants may be delayed or further accelerated by altering the expression of *FvFT1*.

Silencing *FvFT1* partially compensated for the lack of functional FvTFL1, as we observed an increased percentage of inflorescences with two lateral branches on the primary axis and an increased number of branching iterations. In contrast, the ectopic *FvFT1* expression in the *fvtfl1* mutants caused pleiotropic changes in plant architecture by inducing flowering in young seedlings, thus reducing overall plant size, and resulting in inflorescences with only a few flowers. Furthermore, in the functional FvTFL1 background, the overexpression of *FvFT1* resulted in an inflorescence phenotype intermediate between *fvtfl1* and *fvtfl1//FvFT1-OX*, in line with the proposed antagonistic function of these genes.

Recently it was shown that the mutation of *FvLFYa* in woodland strawberry causes the homeotic conversion of flower organs into bracts and the production of additional flowers (Zhang et al., 2022), indicating a conserved function for specifying FM identity. In Arabidopsis and rice, FT and TFL1 proteins are known to competitively bind to bZIP transcription factor FD and 14-3-3 proteins (Zhu et al., 2021; Kaneko-Suzuki et al., 2018). When formed, the FT-FD complex was found to act as a transcriptional activator of *LFY* and *AP1*, whereas TFL1-FD was found to act as a repressor (Zhu et al., 2021). In our experiments, both *FvLFYa* and *FvAP1* were downregulated in the plants with either functional FvTFL1 or silenced *FvFT1* in *fvtfl1* background, suggesting a similar regulatory mechanism.

Altogether, our experimental data indicate that FvTFL1 and FvFT1 antagonistically regulate the complexity of the woodland strawberry inflorescence architecture. With the help of a computational model, we have further shown that the observed diversity of strawberry architectures can be attributed to heterochrony, where FvTFL1 delays, and FvFT1 accelerates the cessation of branching and the production of flowers.

## MATERIALS AND METHODS

### Plant material

Nine *Fragaria vesca* and seven *F. vesca semperflorens* (*fvtfl1* mutant) accessions were used in this study (Supplemental Table S2). Previously reported *FvFT1* overexpression (OX) and RNA silencing (RNAi) lines in *fvtfl1* (*Hawaii-4*) background were used (Koskela et al., 2012; Rantanen et al., 2014). The specificity of the RNAi construct to *FvFT1* was confirmed by analyzing the expression of *FvFT2* and *FvFT3* in the FM tissues (Supplemental Figure S6A – B). To obtain *FvFT1-OX* plants in *FvTFL1* background, we crossed *FvFT1-OX5 (Hawaii-4)* line with *FvTFL1* accession *(FIN56)* and self-pollinated the progeny. F_2_ plants carrying FIN56 *FvTFL1* allele and the desired *FvFT1* construct were used in the experiments.

### Growth conditions and phenotyping

All plants were grown in greenhouse conditions with controlled temperature and photoperiod. In the greenhouse, the plants were illuminated with 150µmol m^-2^ s^-1^ light intensity using high-pressure sodium lamps (Airam 400W, Kerava, Finland). Germinated from seeds or clonally propagated plants were first potted into 7×7cm plastic pots filled with peat moss (Kekkilä, Finland), and kept in the greenhouse under plastic covers for two weeks. Plants were pre-grown at 18°C and an 18h photoperiod for four - six weeks, and periodically supplemented with fertilizer (NPK 17-4-25; Kekkilä). Plants were then potted into 13cm Ø pots and grown until the inflorescences were fully formed. Plants were irrigated with tap water supplemented with fertilizer (NPK 17-4-25; Kekkilä)

For the analysis of FvFT1 and FvTFL1 functions we pre-grew 8 – 13 seed germinated plants per transgenic line or cultivar, or 6 – 13 clonally propagated plants per accession with functional FvTFL1, in growth rooms under an 18h/6h (day/night) photoperiod (AP67, Valoya, Finland) and 25°C temperature. For floral induction, the plants were then moved to a 12h/12h (day/night) photoperiod at 18°C in the greenhouse for six weeks. After the treatment, the plants were kept under 18°C and an 18h/6h (day/night) photoperiod until the inflorescences were fully developed. Inflorescence architecture was analyzed by counting the total number of flowers and flower buds, the number of branching iterations in the longest branching path, and the number of branches on the primary axis. For the functional analysis of FvTFL1, the first fully developed inflorescence was analyzed from each plant. For the functional analysis of FvFT1, five fully developed inflorescences were analyzed.

### Genotyping

DNA was extracted from young, folded leaves of *fvtfl1* mutants and WT plants listed in Supplemental Table S2 as in Koskela et al. (2012). PCR amplified *FvTFL1* fragments were excised from agarose gel, purified, and sequenced using primers listed in Supplemental Table S3.

### RNA extraction and qPCR analysis

Meristems were dissected under stereomicroscope (Zeiss Stemi 2000). SAMs, IMs, and FMs were excised and collected into separate Eppendorf tubes. 5 – 10 meristems were pooled into one tube and used for RNA extraction. Total RNA was extracted from pooled meristem samples as in Mouhu et al. (2009) and treated with rDNase (Macherey-Nagel GmbH, Düren, Germany). 500ng of total RNA was used for cDNA synthesis using ProtoScriptII reverse transcriptase. RT-PCR was performed using a LightCycler 480 SYBR Green I Master kit (Roche Diagnostics, Indianapolis, US) and a Roche LightCycler (Roche Diagnostics, Indianapolis, US) with three technical replicates for each of the tested genes. *FvMSI1 (FvH4_7g08380)* was used as a reference gene. The qPCR primers used in this study are listed in Supplemental Table S3.

### SEM imaging

Dissected meristem samples were fixed in an FAA buffer (3.7 % formaldehyde 5 % acetic acid, and 50 % ethanol) overnight and transferred through ethanol series (50 %, 60%, 70 %, 80 %, 90 %, 100%, 100 %) under mild vacuum (∼0.6 atm). Critical point drying was done using a Leica EM CPD300 (Leica Mikrosystems GmbH, Vienna, Austria). The dried samples were then coated with platinum by a Quorum Q150TS coater (Quorum Technologies, UK). The samples were examined under a Quanta 250 FEG (FEI, Hillsboro, OR, USA) scanning electron microscope located at the Electron Microscopy Unit (Institute of Biotechnology, University of Helsinki). Pseudo-coloring was done in Adobe Photoshop CC 2019.

### Statistical analyses

The phenotypic data (flowering time, number of flowers per inflorescence and branching iterations) were analyzed using random intercept models, *y* = *β*_0_ + *β*_1_*X* + *Gt* + *ε*, where *y* = dependent variable, *β*s = fixed effects, *X* = design matrices for *FvTFL1* background or *FvFT1* transgenic construct, *Gt* = random accession or independent line effect, and *ε* = an error term. Poisson distribution model was applied for count data. Statistical analyses were performed using the R/lme4 package (Bates et al., 2015). All analyses were performed in R version 4.1.1 (R Core Team 2021).

### Inflorescence modelling

The models were written in the L-system-based L+C plant modeling language (Karwowski and Prusinkiewicz, 2003) and executed using the lpfg simulator incorporated into the Virtual Laboratory (vlab) v5.0 plant modeling software (algorithmicbotany.org/virtual_laboratory). All simulations were performed on MacBook Pro computers under macOS High Sierra 10.13.6. The supplemental materials include three complete vlab objects (versions of the model): the basic version with a simplified representation of plant organs (bracts and flowers), convenient for analyzing the model logic; the extended version with full representation of organs, used in the model calibration shown in Figure 7; and the same extended version configured to simulate the diverse phenotypes shown in Figure 8. The key elements of the model code are discussed in the Supplemental Model Description. Parameters used to simulate the diverse phenotypes were found by interactively exploring the model parameter space using the tools included in vlab (control panels and graphically defined timelines and functions). The reference model (Figure 7) was calibrated using the method described by Cieslak et al. (2022).

## Supporting information

Supplemental model description

## Acknowledgements

We thank Elli Koskela for providing seeds; Eija Takala and Marjo Kilpinen for laboratory assistance; Katriina Palm for maintaining the plant materials; the staff of the Electron Microscopy Unit at the Institute of Biotechnology, University of Helsinki, for assistance. This work was supported by Jane and Aatos Erkko Foundation grant 4709012 (to P.E. and T.H.), the Academy of Finland grant 317306 (to T.H), Natural Sciences and Engineering Research Council of Canada Discovery Grant 2019-06279 (to P.P.), the Plant Phenotyping and Imaging Research Centre/Canada First Research Excellence Fund (to P.P. and M.C.), and the Finnish Cultural Foundation grant 00221168 (to S.L.). S.L. was affiliated with the Doctoral Programme in Plant Sciences (DPPS, University of Helsinki).

## Footnotes

* The concept of Petri nets formally underlying Figure 6B is further described by Peterson (1981), and in application to plant modeling by Prusinkiewicz and Remphrey (2000). The use of Petri nets enhances the flowchart representation of inflorescence development proposed by Kellogg (2000) by explicitly representing events that produce multiple organs, which then develop concurrently.

