## Supplemental model description for "Diversity of the woodland strawberry inflorescences results from heterochrony antagonistically regulated by *FvTFL1* and *FvFT1*"

### 1. General information

We have implemented three versions of the model:

The `Strawberry-basic` version, focused on in this Description, captures the essence of the development of the branching structure of wild strawberry inflorescences. Flowers and fruits are represented schematically, as colored spheres. The gradual unfolding of young bracts is not simulated. The model generates the architecture of all inflorescences shown in Fig. 7 (Phenotype 0) and Fig. 8 (Phenotypes 1–7), depending on the selected parameter values.

The `Strawberry-calibrating` version augments the basic model with a more realistic representation of organs. It illustrates model calibration according to a developmental sequence of photographs of a real inflorescence (Fig. 7).

The `Strawberry-diversity` version employs the same representation of organs as `Strawberry-calibrating`, but has been configured to generate the diverse phenotypes shown in Fig. 8 by changing the parameter values.

All models were specified in the L-system-based L+C modeling language [Karwowski, 2002, Karwowski and Prusinkiewicz, 2003]; for a tutorial introduction see [Prusinkiewicz et al., 2018] and for further examples focused on model calibration see [Cieslak et al., 2022]. Simulations were effected using the lpfg simulator included in the Virtual Laboratory plant modeling platform (vlab) available at `http://algorithmicbotany.org/virtual_laboratory/`. Vlab manuals provide additional documentation of the L+C language and auxiliary vlab tools (control panels, timeline editors, function and surface editors) needed to manipulate and calibrate the model interactively. The complete models (vlab objects) are provided as supplementary materials; in the description below we focus on the essential element, the simulation of the branching architecture of the strawberry inflorescence.

### 2. Vegetativeness calculation

The simulation is driven by the progress of time $t$ and the gradual decrease of $veg$ represented by an exponential function of time,

$$veg(t) = veg_{init} e^{-rt},$$

where $veg_{init}$ is the vegetativeness value at the beginning of the simulation and $r$ is the decline rate (cf. Fig. 6A). As the simulation progresses over discrete time intervals, the *veg* value is decremented according to a differentiated version of the above equation,

$$\frac{dveg}{dt} = -veg_{init} r e^{-rt},$$

which is approximated as

$$veg(t+dt) \approx veg(t)(1-rdt).$$

In the last equation, $dt$ is a small, but not infinitesimal, time increment. The corresponding implementation in the code (line 169) is:

```
169    veg -= veg*veg_decline_rate*dt;
```

Instead of specifying the decline rate $r$ directly, we have found it more intuitive to specify time $t_{1/2}$ at which *veg* decreases to one half of its initial value. The decline rate and the half-life time are related by the equation

$$\frac{1}{2} veg_{init} = veg_{init} e^{-rt_{1/2}}$$

and thus

$$r = \frac{\ln(2)}{t_{1/2}},$$

which is implemented as

```
   veg_decline_rate = log(2) / veg_half_life;
```

### 3. Code structure

The basic module type on which the model operates is a meristem, defined as follows:

```
struct MeristemData {
   int state;  // MONOPODIAL, SYMPODIAL, or FLOWERING
   float age;  // age since creation or production of the most recent primordium
   float d;    // diameter
};
module Meristem(MeristemData);
```

According to this definition, a meristem is characterized by three variables. The `state` can assume values `MONOPODIAL`, `SYMPODIAL` or `FLOWERING` (see Petri net in Fig. 6B). The `age` measures time since the production of the previous primordium. This information is needed to produce branches at prescribed time intervals (plastochrons). The value `d` determines the diameter of the internode that supports the meristem.

As the simulation progresses, the meristem age and diameter value are updated by the rule (production):

```
Meristem(m):
{
   m.age += dt;
   m.d = fmax(m.d, distal_diameter * meristem_size(m.age));
   produce Meristem(m);
}
```

where `meristem_size()` is a graphically defined function of `age`, increasing meristem diameter from 0 to `distal_diameter` over time. The `fmax()` function assures that diameter `d` never decreases, thus preventing fluctuations of the internode width as the meristem produces consecutive primordia.

Bract data are updated more simply:

```
Bract(b):
{
   b.age += dt;
   b.d = distal_diameter * petiole_width(b.age);
   produce Bract(b);
}
```

Beginning with meristems and bracts, the diameter values propagate basipetally, increasing at each node (branching point) according to the equation [Shinozaki et al., 1964, MacDonald, 1983]:

$$d_p^n = \sum_c d_c^n,$$

where $d_p$ is the diameter of the parent internode, each term $d_c$ is the diameter of an organ (meristem or bract) or a child internode supported by this parent, and the exponent $n$ is a model parameter. Our implementation of the pipe model (lines 220-240 of the code, not quoted in this Description) is based on [Karwowski and Prusinkiewicz, 2003].

The production of new primordia and changes in the meristem state are captured by two further rules (formally, these are decomposition rules explained, for example, in [Karwowski and Prusinkiewicz, 2003]). The first rule has the following structure:

```
Meristem(m):
{
   if (m.age >= plastochron) {
      // Code for handling monopodial meristem
      ...
      // Code for handling sympodial meristem
      ...
   }
}
```

As specified in the code, this rule only takes effect when the local `age` of a meristem has reached the `plastochron` time. Further action then depends on the meristem state (Fig. 6B). The portion of the code applying to the meristem in the `MONOPODIAL` state is as follows:

```
253       if (m.state == MONOPODIAL) {  // 1st order meristem: monopodial branching
254          if (veg > th_m) {          // high veg: produce a lateral meristem
255             m.age -= plastochron;
256             InternodeData s = {m.age, veg, 0};
257             nproduce SB Left(angle_lateral) Primordium(m.age-delay_mono, veg-th_s) EB;
258             produce Right(angle_main) RollL(180) Internode(s) Meristem(m);
259          }
260          else {
261             m.state = FLOWERING;    // low veg: change state to flowering
262             produce Meristem(m);
263          }
264       }
```

If `veg > th_m`, the meristem produces a lateral primordium (line 257) and recreates itself (line 258) with the age reduced by `plastochron` (line 255). Consistent with the Petri net Event 1, the recreated meristem is supported by a new internode (initialized in line 256). Important attributes of this production are the `delay_mono` parameter, which defines the difference in the development of the lateral primordium with respect to the main meristem; `angle_lateral`, which determines the branching angle that the axis of the new primordium forms with respect to the main axis; and `angle_main`, which determines the deviation of the main apex from its previous course following the production of the lateral primordium. The `RollL(180)` module in line 258 defines the phyllotaxis of the main axis as opposite; this is not consistent with the actual architecture of the strawberry inflorescence, but facilitates visual comparisons of the model with photographs, in which the inflorescence was laid out flat on the supporting surface to better expose its structure. Further attributes of the primordium are captured by a separate decomposition rule, discussed below. Alternatively (i.e., if `veg` ≤ `th_m`), the main meristem transitions to the `FLOWERING` state (line 261), thus implementing Petri net Event 2.

The portion applying to the meristem in the `SYMPODIAL` state has the form:

```
266      if (m.state == SYMPODIAL)  {  // meristem of order 2 or higher: sympodial branching
267         m.age -= plastochron;
268         InternodeData s = {m.age, veg, 0};
269         nproduce SB Left(angle_sympodial) Primordium(m.age-delay_sym, veg-th_s) EB;
270         nproduce SB RollL(180) Left(angle_sympodial)
271            Primordium(m.age-delay_sym-delay_diff, veg-th_s-th_diff) EB;
272         m.state = FLOWERING;       // parent meristem always switches to flowering
273         produce Internode(s) Meristem(m);
274      }
```

Consistent with Petri net Event 4, a sympodial meristem produces two new lateral primordia (lines 269-271), and switches to the `FLOWERING` state (line 272); a supporting internode is also produced (line 273). Parameters of this process are: `delay_sym`, which describes the difference in the developmental stage of the lateral primordia compared to the meristem that produced them in a manner analogous to `delay_mono`; `delay_diff`, which can capture a possible difference in the developmental stage of lateral primordia, and `th_diff`, which introduces a similar difference in the `th_s` threshold value determining the fate of these primordia. Finally, the `angle_sympodial` parameter determines the branching angle between the axes of the lateral primordia and that of their supporting internode.

The second rule pertinent to branching specifies the fate of primordia:

```
278 Primordium(p_age,p_veg) :
279 {
280    BractData b = {p_age, veg, (float) (bract_size(b.age))};
281    if (p_veg > 0) {                            // If veg is high enough
282       MeristemData new_m = {SYMPODIAL, p_age};
283       InternodeData s = {new_m.age, veg, 0};
284       nproduce SB Left(bract_angle_branch) Bract(b) EB;  // produce a bract
285       produce Internode(s) Meristem(new_m);    // and a meristem supported by an internode
286    }
287    else {                                      // If veg is not high enough
288       nproduce SB Right(bract_angle_no_branch) Bract(b) EB; // produce a bract
289       produce;                                 // and abort
290    }
291 }
```

The primordium has two parameters. The `p_age` parameter defines the initial age of the primordium. This age may be different from the age of the parent meristem, as defined by the paramaters `delay_mono`, `delay_sym` and `delay_diff` discussed above in the context of the `Meristem` rule. The second parameter, `p_veg`, describes local vegetativeness of the primordium at the time of its creation. This value is reduced from the globally defined `veg` by threshold `th_s` and, possibly, `th_diff` (line 271); consequently, the Petri net condition $veg > th_s$ distinguishing between Events 3 and 5 (Fig. 6B) is implemented as `p_veg > 0` in the `Primordium` rule. If this condition is satisfied, the rule implements Event 3 by decomposing a primordium into a bract and a lateral meristem — supported by an internode — in the bract's axil (lines 281-286). In the opposite case, the rule implements Event 5 by only yielding a bract (lines 287-290). With a non-zero `th_diff`, the two lateral primordia produced by a cymose inflorescence may have different fates, such that one primordium yields a lateral meristem, and another one does not.

### 4. Parameter space exploration

The repetitive application of the production advancing the age of primordia, followed periodically with the decomposition rules for meristems and primordia, creates the compound thyrse architecture characteristic of strawberry inflorescences. The extent of this structure — the number of sympodial branches supported by the primary axis and the number of repetitions of cympose branching — depends on two factors. The first factor is the timing of the initiation of new primordia, controlled directly by the `plastochron` (set in the model to one unit of time) and delay values `delay_sym` and `delay_diff`. The second factor is the timing of the switches in meristem state, controlled by the relation between `veg` and thresholds `th_m` and `th_s` (Fig. 6A), and possibly adjusted by the `th_diff` value. Returning to mathematical notation, time $t_m$ at which condition $veg < th_m$ triggers the production of the flower terminating the monopodial axis (Petri net Event 2) can be expressed as

$$veg_{init}e^{-rt_m} \approx th_m,$$

which is equivalent to

$$t_m \approx \frac{1}{r}(\ln(veg_{init}) - \ln(th_m)).$$

(Aproximations are due to the fact that *veg* values in the model are only sampled when the meristem age exceeds the plastochron, which may introduce some delay.) Likewise, time $t_s$ at which condition $veg < th_s$ arrests the production of sympodial meristems can be expressed as

$$t_s \approx \frac{1}{r}(\ln(veg_{init}) - \ln(th_s)).$$

From these equalities it follows that different combinations of values $veg_{init}$, $r$, $th_m$, and $th_s$ may produce the same switch times $t_m$ and $t_s$. When exploring the model, we have focused on the *veg* decline rate $r$, manipulated through the half life of *veg*, $t_{1/2}$, as discussed earlier. This choice has the appeal of simplicity, with a single parameter $t_{1/2}$ controlling both switch times as well as the appearance of consecutively produced internodes and bracts in a coordinated fashion. Moreover, the action of genes *FvTFL1* and *FvFT1* can be interpreted in a straightforward manner as factors decreasing or increasing $t_{1/2}$. Nevertheless, the action of genes can also be mapped on other model parameters (for instance, the initial vegetativeness, $veg_{init}$) with similar effect. We have thus shown that the diversity of wild strawberry inflorescence architectures can be captured by a simple model, but more molecular-level data are needed to fully resolve how *FvTFL1* and *FvFT1* control model parameters.
