## Supplemental figures and tables for "Diversity of the woodland strawberry inflorescences results from heterochrony antagonistically regulated by *FvTFL1* and *FvFT1*"

**Supplemental Table S1.** Model parameters used to produce inflorescences in Figure 7 (Phenotype 0) and Figure 8 (Phenotypes 1-7). Parameters shown in bold are discussed in the main text or Supplemental Model Description; the remaining parameters are explained in the code. Only values differing from the previous column are shown. The key parameter differentiating the models is `veg_half_life`, which determines the time (measured in plastochrons) over which *veg* decreases to ½ of its initial value. Related parameters are grouped together as in the code.

| **Phenotype** | **0** | **1** | **2** | **3** | **4** | **5** | **6** | **7** |
|---|---|---|---|---|---|---|---|---|
| max_age | 8 | | | | | | | |
| **dt** | 0.02 | | | | | | | |
| **plastochron** | 1 | | | | | | | |
| **delay_mono** | 0.44 | | | | | | | |
| **delay_sym** | 0.3 | 0 | | | | | | |
| **delay_diff** | | | | | | | | |
| **veg_init** | 1 | | | | | | | |
| **veg_half_life** | 1.98 | 2.77 | 2.42 | 2.07 | 1.64 | 1.37 | 1.05 | 1.05 |
| **th_m** | 0.43 | | | | | | | |
| **th_s** | 0.27 | | | | 0.23 | | | 0.6 |
| **th_diff** | 0.08 | | | | 0 | | | |
| b_th_1 | 0.6 | | | | | | 0.48 | |
| b_th_2 | 0.45 | | | | | | | |
| pleiotropic_scale | 4 | 1 | | | | 0.82 | 0.5 | 0.4 |
| peduncle_length_ctrl | 0.4 | 0.2 | | 0.3 | 0.65 | 0.7 | 0.75 | 0.6 |
| int_len_power | 1.2 | | | | | | | |
| internode_segments | 10 | | | | | | | |
| **distal_diameter** | 0.011 | | | | 0.008 | 0.011 | | |
| **pipe_model_power** | 3 | | | | | | | |
| **angle_main** | 12 | 35 | 38 | 17 | 27 | | | 35 |
| **angle_lateral** | 10 | 17 | 13 | 29 | 7 | | | 17 |
| **angle_sympodial** | 20 | 32 | | | 35 | | 32 | |
| int_elasticity | 0.32 | 0.06 | | | | | | |
| int_elasticity_power | 1.3 | | | | | | | |
| bract_angle_branch | 5 | | | | | | | |
| bract_angle_no_branch | 25 | | | | | | | 17 |
| ang_adjustment_1 | 0 | | | | | | 6 | 14 |
| ang_adjustment_2 | 35 | | | | | | | |
| petiole_length_1 | 0.2 | 0.7 | | | | | 0.3 | 0.3 |
| petiole_length_2 | 0.02 | | | | | | | |
| petiole_elasticity_1 | 1.8 | | | | | | | |
| petiole_elasticity_2 | 1 | | | | | | | |
| bract_size_power | 0.5 | | | | | | | |

**Supplemental Table S2.** Plant material used in this study.

| Accession Name | Origin/Collection Site |
|---|---|
| *Alexandria* (*PI602923*) | NCGR* |
| *Baranovka* | Lyubov Kuznetsova** |
| *Baron Solemacher* (*PI551507*) | NCGR* |
| *Hawaii-4* (*PI551572*) | NCGR* |
| *Piikkiö* | Unknown |
| *Reine des Vallees* (*PI551824*) | Dr. Amparo Monfort*** |
| *Yellow Wonder* (*PI551827*) | NCGR* |
| *CRO2* | 44º46’15.1896’’ (N) 15º38’51.6408’’ (E) |
| *FIN56* (*PI551792*) | NCGR* |
| *FR4* | 48º25’24.4452’’ (N) 7º39’48.0168’’ (E) |
| *GER4* | 50º59’6.9036’’ (N) 11º19’21.216’’(E) |
| *GER10* | 47º33’21.168’’ (N) 10º1’17.2632’’ (E) |
| *ICE10* | 64º45’36.774’’ (N) -21º35’35.2824’’ (E) |
| *IT1* | 45º56’13.362’’ (N) 10º48’20.2392’’ (E) |
| *IT7* | 46º16’30.1188’’ (N) 11º16’54.9372’’ (E) |
| *LIT1* | 54º47’51.7812’’ (N) 25º20’54.2292’’ (E) |

*National Clonal Germplasm Repository, Corvallis, USA; ** The Federal Research Center Institute of Cytology and Genetics, Novosibirsk, Russia; *** Centre for Research in Agricultural Genomics, Barcelona, Spain

**Supplemental Table S3.** Primers used in this study.

| Name | Forward | Reverse | Reference |
|---|---|---|---|
| *FvTFL1 (FvH4_6g18480) (cloning/sequencing)* | ATGGCAAGAATGTCGGAACC | CTAGCGTCTTCTTGCTGCC | |
| *FvTFL1 (FvH4_6g18480) (qPCR)* | AACGGCAGCAACAGGAAC | CTGGCACCACAGATGCTACA | Koskela et al., 2012 |
| *FvAP1 (FvH4_4g29600) (qPCR)* | AGCTCAGGAGGTTCATGACTG | TAAGGTCGAGCTGGTTCCTC | Koskela et al., 2012 |
| *FvMSI1 (FvH4_7g08380) (qPCR)* | TCTCCACACCTTTGATTGCCA | ACACCATCAGTCTCCTGCCAAG | Mouhu et al., 2009 |
| *FvFT1 (FvH4_6g00090) (qPCR)* | CAATCTCTTGGCCGAAAACT | TGAGCTCAAACCTTCCCAAG | Koskela et al., 2012 |
| *FvLFYa (FvH4_5g09660) (qPCR)* | GATGACAGAATCAATGGAGGAG | CTGGTTTGTGACCTTGGTGG | |
| *FvFT2 (FvH4_4g30710) (qPCR)* | ACTCGGTGGCTTGTGTTTTC | ATCACTCTCCCGACGACAAG | Nakano et al., 2015 |
| *FvFT3 (FvH4_3g09870) (qPCR)* | AGCCGTTCACCAAGTCTGTG | GTGGACAACATGAGAAGGTTTG | Nakano et al., 2015 |

### Figures

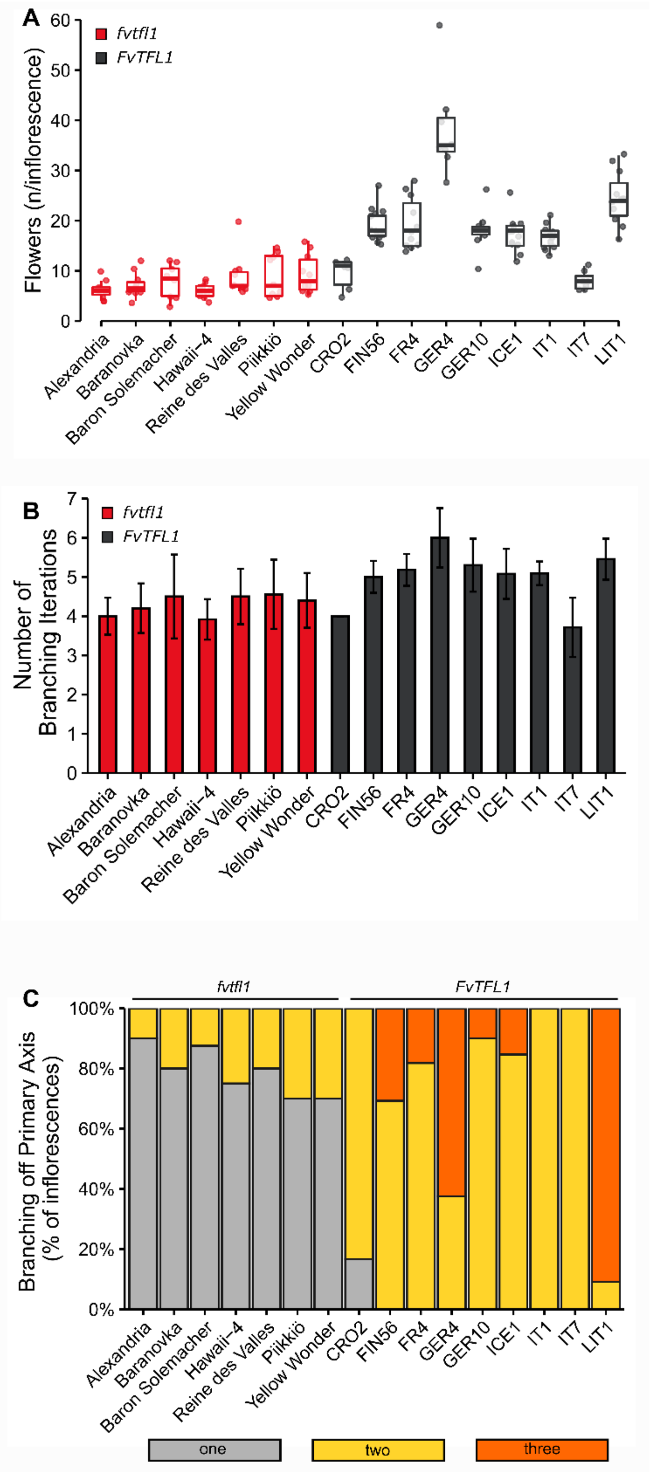

**Supplemental Figure S1. Inflorescence phenotypes of seven *fvtfl1* and nine *FvTFL1* genotypes. A.** Number of flowers on the first emerged fully-developed inflorescence. Boxplots and points show distribution of raw data. Each point represents an individual inflorescence. **B.** Average number of branching iterations along the longest branching path. Bars and whiskers represent the mean ± standard deviation (n = 6 – 13). **C.** Percentage of observed inflorescences with one, two, or three branches on the primary branching axis.

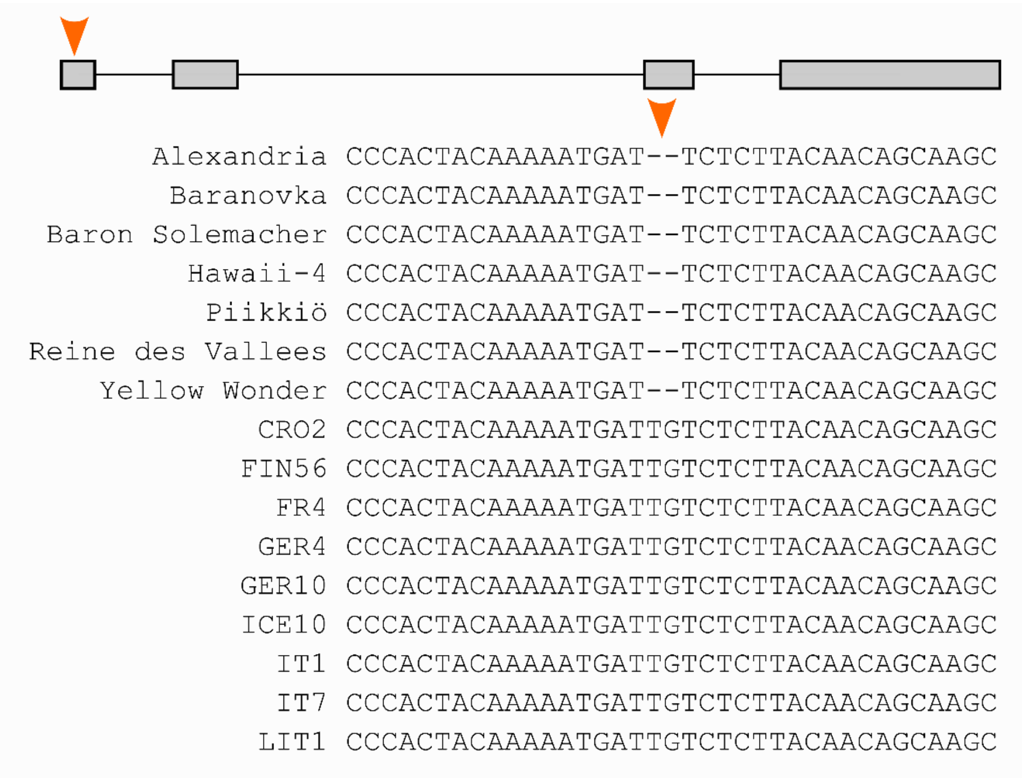

**Supplemental Figure S2.** Alignment of the part of the first exon of *FvTFL1* gene in seven cultivars and nine WT accessions.

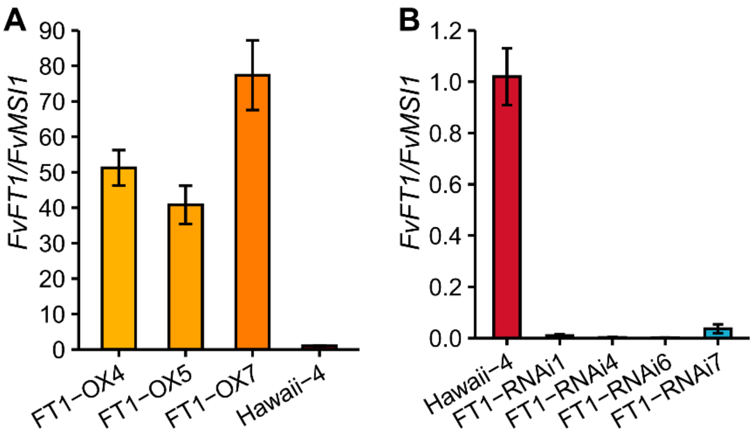

**Supplemental Figure S3.** *FvFT1* expression levels in the leaves of *Hawaii-4* (control), *FvFT1* overexpression (OX) and *FvFT1* silencing (RNAi) transgenic lines. Bars and whiskers show mean ± standard error (n = 3 – 4). Expression levels were normalized to *Hawaii-4*. *FvMSI1* was used as a reference gene.

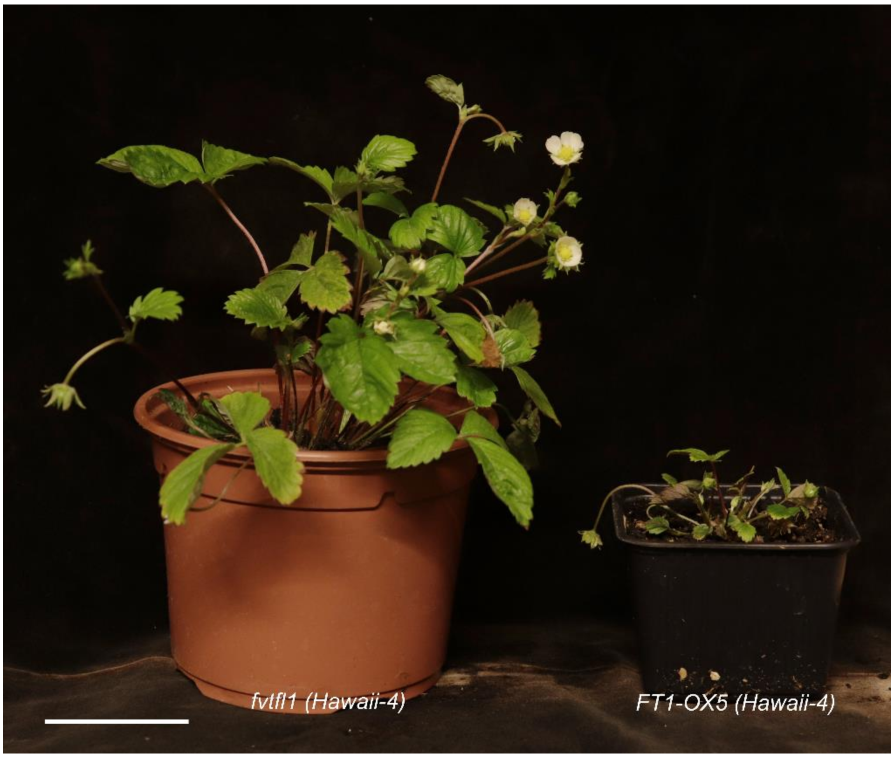

**Supplemental Figure S4.** *FvFT1* overexpression (35S) in the *fvtfl1 (Hawaii-4)* background induces flowering in all axillary buds and reduces overall plant size. Scale bar equals 5 cm.

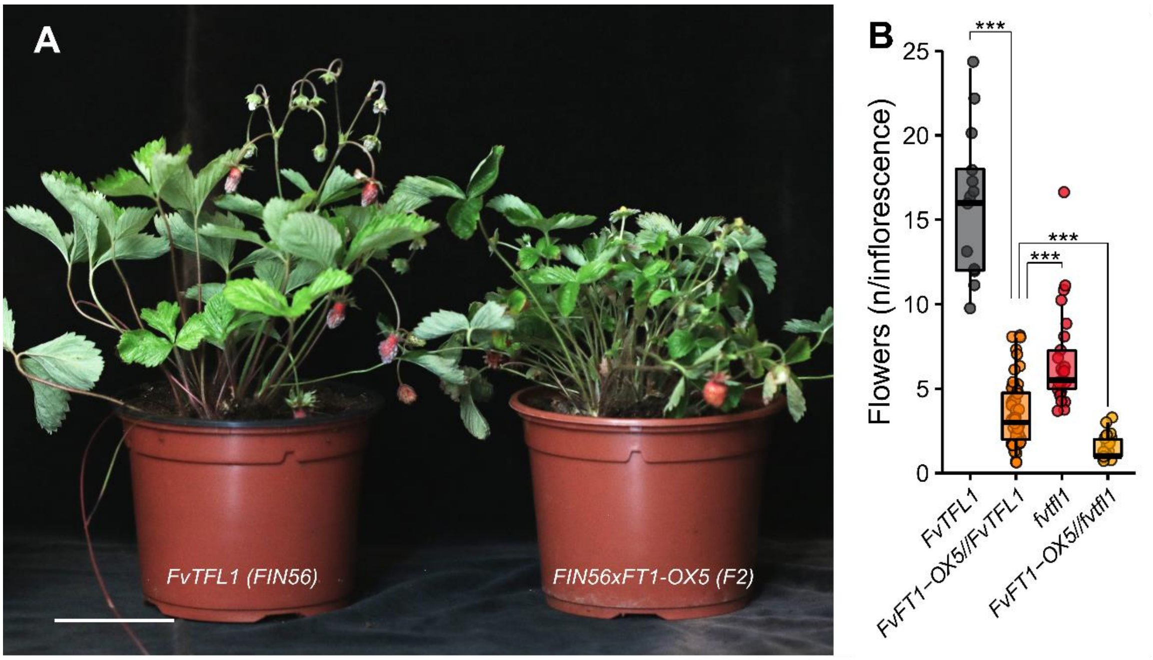

**Supplemental Figure S5.** *FvFT1* Overexpression in the functional *FvTFL1* background. **A.** overall plant phenotypes. **B.** The number of flowers per inflorescence in *FvFT1* overexpression plants in functional and non-functional *FvTFL1* backgrounds.
Boxplots and points show the distribution of raw data. Each point represents an individual inflorescence. Up to six inflorescences per plant were examined. Data were analyzed by fitting a generalized linear model (GLM). *** indicate P value < 0.0001 between the groups (*t*-test).

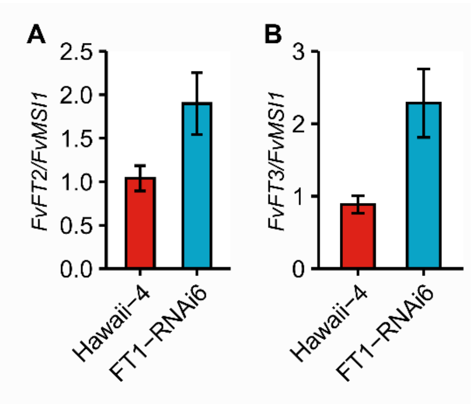

**Supplemental Figure S6.** The expression of *FvFT2* **(A)** and *FvFT3* **(B)** in the FMs of *Hawaii-4* and *FT1-RNAi6* plants. Bars and whiskers show mean ± standard error (n = 3 – 4). Expression levels were normalized to *Hawaii-4.*
